# Vps54 regulates lifespan and locomotor behavior in adult *Drosophila melanogaster*

**DOI:** 10.1101/2021.08.20.457133

**Authors:** Emily C. Wilkinson, Emily L. Starke, Scott A. Barbee

## Abstract

Vps54 is an integral subunit of the Golgi-associated retrograde protein (GARP) complex, which is involved in tethering endosome-derived vesicles to the *trans*-Golgi network (TGN). A destabilizing missense mutation in *Vps54* causes the age-progressive motor neuron (MN) degeneration, muscle weakness, and muscle atrophy observed in the wobbler mouse, an established animal model for human MN disease. It is currently unclear how the disruption of Vps54, and thereby the GARP complex, leads to MN and muscle phenotypes. To develop a new tool to address this question, we have created an analogous model in *Drosophila* by generating novel loss-of-function alleles of the fly *Vps54* ortholog (*scattered/scat*). We find that null *scat* mutant adults are viable but have a significantly shortened lifespan. Like phenotypes observed in the wobbler mouse, we show that *scat* mutant adults are male sterile and have significantly reduced body size and muscle area. Moreover, we demonstrate that *scat* mutant adults have significant age-progressive defects in locomotor function. Interestingly, we see sexually dimorphic effects, with *scat* mutant adult females exhibiting significantly stronger phenotypes. Finally, we show that *scat* interacts genetically with *rab11* in MNs to control age-progressive muscle atrophy in adults. Together, these data suggest that *scat* mutant flies share mutant phenotypes with the wobbler mouse and may serve as a new genetic model system to study the cellular and molecular mechanisms underlying MN disease.

## INTRODUCTION

Neurodegenerative diseases are severe and often fatal disorders associated with reduced function, or loss of function, of neurological components. This degeneration commonly leads to cognitive impairment and/or motor dysfunction. The primary risk factor associated with neurodegeneration is aging, and as a great portion of the population continues to age the prevalence of such disorders continues to increase (Niccoli and Partridge, 2012). The identification of mutations linked to human neurodegenerative diseases have highlighted several important intracellular pathways that are involved in disease pathogenesis. Many of these genes can be categorized by their contribution to critical intracellular processes including RNA and protein metabolism, axonal and cytoskeletal dynamics, and membrane trafficking (Taylor et al., 2016).

Endocytic trafficking has been implicated in several specialized processes in neurons including axon guidance and outgrowth, synaptic plasticity, and axonal transport (Wojnacki and Galli, 2016). Disruption of pathways involved in the function of endocytic trafficking has been linked to progressive neurodegenerative disorders such as amyotrophic lateral sclerosis (ALS), Parkinson’s disease (PD), and hereditary spastic paraplegias (HSPs) (Schreij et al., 2016). MN axons appear to be particularly sensitive to mutations in genes involved in membrane trafficking, specifically ALS and HSPs. The membrane trafficking genes that have been implicated in ALS include *Alsin (ALS2), C9ORF72*, and *Optineurin (OPTN)* (Devon et al., 2006; Hadano et al., 2001; Maruyama et al., 2010; Stepto et al., 2014; Waite et al., 2014; Yang et al., 2001). Genes involved in HSPs are *Spastin (SPG4), Strumpellin (SPG8), Spatacsin (SPG11), Spastizin (SPG15), AP5 (SPG48)*, and *Vps37A (SPG53)* (Hanein et al., 2008; Hazan et al., 1999; Patel et al., 2002; Slabicki et al., 2010; Valdmanis et al., 2007; Zivony-Elboum et al., 2012).

A destabilizing missense mutation in the gene encoding for the vacuolar protein sorting-associated protein 54 (Vps54) causes age-progressive MN degeneration in mice. This mouse model, known as the “wobbler” mouse, is used to model human MN disease because it shares many striking phenotypic similarities with ALS (Moser et al., 2013). Vps54 is a core subunit of the heterotetrametric Golgi-associated retrograde protein (GARP) complex and is involved in tethering retrograde transport carriers, derived from endosomes to the *trans*-Golgi network (TGN) (Bonifacino and Hierro, 2011). The subunits that compose the primary structure of the GARP complex are Vps51, Vps52, Vps53, and Vps54 (Conibear and Stevens, 2000). Destabilization of Vps54 in the wobbler mouse leads to a compensatory decrease in levels of Vps53 and disruption of the assembly of the GARP complex (Perez-Victoria et al., 2010). The N-terminus of yeast and mammalian Vps54 binds to TGN-associated soluble N-ethylmaleimide-sensitive fusion protein attachment protein receptors (t-SNAREs) while the C-terminus interacts with endosomes (Perez-Victoria and Bonifacino, 2009; Quenneville et al., 2006). Knockdown of Vps54 and other GARP complex subunits results in defects in retrograde and anterograde vesicle transport (Conibear and Stevens, 2000; Hirata et al., 2015; Perez-Victoria and Bonifacino, 2009; Perez-Victoria et al., 2008). Additionally, knockdown of GARP complex subunits causes lysosomal dysfunction (Perez-Victoria and Bonifacino, 2009; Perez-Victoria *et al.*, 2008). Taken together, these data strongly suggest that Vps54 (and the GARP complex) plays a conserved and essential role in endolysosomal trafficking pathways.

*Drosophila melanogaster* have a single ortholog of Vps54 called *scattered* or *scat*. We have previously shown that disruption of *scat* causes defects in the development of the *Drosophila* larval neuromuscular junction (NMJ) (Patel et al., 2020). Moreover, we found that presynaptic *scat* interacts genetically with *rab7* to regulate the composition of the postsynaptic density via an unknown trans-synaptic mechanism (Patel *et al.*, 2020). We hypothesized that these changes at the larval NMJ may precede neurodegenerative phenotypes in aging adults. Here we demonstrate that loss of *scat* expression leads to a significant reduction in adult lifespan. We show that *scat* mutants have sex-specific defects in lifespan, body size, and muscle mass with females exhibiting a more severe phenotype. Female *scat* mutants also exhibit neurological dysfunction (seizure) and age-progressive defects in locomotor behavior. Finally, we demonstrate that the simultaneous MN-specific disruption of *scat* expression and *rab11* function exacerbates muscle atrophy in adult females, suggesting phenotypes are due to a trafficking defect. These data suggest that the *scat* loss-of-function model shares many phenotypes with the wobbler mouse, making it a useful tool to study the mechanisms underlying MN disease.

## MATERIALS AND METHODS

### *Drosophila* lines and genetics

The following fly lines were obtained from the Bloomington *Drosophila* Stock Center: *w^1118^*, *scat^1^cn^1^, C380-Gal4, UAS-Scat^TRiP^ (HMS01910), UAS-LUC.VALIUM10, UAS-YFP:Rab5, UAS-YFP:Rab5 (S43N)*, *UAS-YFP:Rab7, UAS-YFP:Rab7(T22N), UAS-YFP:Rab11, UAS-YFP:Rab11(S25N*). In our hands, the *cn^1^* allele caused significant differences in locomotor assays relative to controls (our unpublished observations). Therefore, *cn^1^* was recombined away from new *scat* alleles. Three new fly lines were generated by mobilizing the transposable element insertion in the *scat^1^* allele by introducing the Δ2-3 transposase into the genome following standard procedures. The two resulting deletion lines (*scat*^Δ*244*^ and *scat*^Δ*312*^) and one precise excision line (*scat^329PE^)* were screened by PCR and DNA sequencing. All fly crosses were maintained on standard Bloomington media at 25°C, 65% humidity, and a 12:12 hour light-dark cycle. Unless otherwise noted, *w^1118^* (*Iso31*) was used as the wild-type control. For overexpression and short hairpin RNAi studies, the UAS/GAL4 system was used (Brand and Perrimon, 1993). To co-overexpress transgenes in MNs, single copies of all indicated elements were crossed into a background containing one copy of the GAL4 transgene. The GAL4 line used in this work was the MN-specific driver, *C380-GAL4* (Budnik et al., 1996).

### Longevity assay

Male and female flies from each genotype were collected within 24 hours of eclosion and segregated by sex. Populations of 300 flies were used in each cohort. All flies were transferred onto fresh food every 2 days and scored for survival at transfer.

### Determination of gender ratios, eclosion, and quantification of body size

For the quantification of gender ratios, 100 adult flies from each genotype were collected at random over a 24-hour period post-eclosion and then sexed. For the determination of eclosion percentages, 50 wandering third instar larvae of each genotype were collected, allowed to pupate, and adult flies collected and sexed. For the analysis of adult body size, 5 adults of each sex and genotype were collected within 24 of eclosion and allowed to age for 24 hours. Flies were anesthetized with CO_2_ and the ventral side of the abdomen was imaged using Leica S9i stereo microscope with 10MP CMOS-camera. Length was determined by drawing and measuring a line from the rostral to caudal ends using the measurement tools in open-source Fiji/ImageJ2. Size was determined by drawing a line around the thorax and abdomen and calculating area.

### Paraffin embedding, sectioning, and image analysis

Flies were collected within 24 hours of eclosion and allowed to age until the desired time point. Flies were anesthetized using CO_2_ and oriented in a custom 3D printed embedding collar so that the thorax was oriented towards the blades. Flies in the collar were then incubated overnight at 4°C with Carnoy’s fixative. Flies were then dehydrated by sequentially incubating for 20 minutes each in room temperature 40%, 70%, and 100% ethanol. Flies were then transferred to a 1:1 solution of methyl benzoate: paraffin wax and incubated for 1 hour at 65°C. For embedding, flies in the collar were transferred to a foil pocket which was then filled with melted paraffin wax and incubated at 65°C for 2 hours. Following incubation, the pocket was stored overnight at room temperature to allow the wax to harden. Paraffin embedded tissue was sectioned into 10 μm sections using a Leica RM2125 microtome and floated on cold water. Sections were collected using charged glass microscope slides and allowed to dry for 2 hours at room temperature. Slides were deparaffinized by incubation in room temperature xylene for 15 minutes. Tissue was rehydrated prior to staining by sequentially incubating for 10 minutes each in 100%, 95%, 80% ethanol and ddH_2_O. Tissue was stained by incubating slides with hematoxylin for 5 minutes followed by eosin for 30 seconds with to washes in between treatments (diH_2_O followed by 95% ethanol). Tissue was then dehydrated by treating incubating slides for 15 minutes each in 95% then 100% ethanol and stored in xylene overnight at room temperature. Permount, a xylene based mounting media, was used to mount and store stained sections. Sections were imaged using a Laxco SeBa 2 series digital microscope system with a 10X objective (N.A. 1.25). Image analysis was done using the measurement tools in Fiji/ImageJ2. Muscle area was determined by tracing around the perimeter of all 12 dorsal longitudinal muscles together excluding any space between individual muscles. Thoracic area was determined by tracing around the cuticle. Sections with less than ~ 90% intact cuticle were excluded from analysis.

### Spontaneous flight assay

For the analysis of flight behavior, 20 female flies of each genotype were collected within 24 hours of eclosion and allowed to age until the desired time point and briefly stored in an empty containment vial. Flies were lightly tapped to the bottom of the vial and then dropped from a unform height into a funnel on top of a 500 ml graduated cylinder. The inside of the cylinder was coated with a thin layer of mineral oil. When flies enter a free fall, they will attempt to fly to and land on the nearest surface. When flies land on the side of the cylinder they become stuck in the mineral oil. Flies with poor locomotor function are expected to fall farther in the cylinder before landing on the side. The height of the graduated cylinder was separated into equal quadrants. The quadrant at which each fly landed and stuck to the side of the cylinder was recorded for each genotype.

### Negative geotaxis assay

For the analysis of climbing behavior, 20 female flies of each genotype were collected within 24 hours of eclosion and allowed to age until the desired time point. Flies tapped to the bottom of a container exhibit a climbing reflex, where they favor climbing over flight to regain position at the top of a container. Flies were transferred to an empty vial marked with the target height, gently tapped down to the bottom, and allowed to climb up the sides for 90 seconds. Because *scat* mutant flies were generally poor climbers, the target height was set relatively low (1 cm). Data was recorded using the built-in camera on a MacBook Air and Photo Booth video recording software. All genotypes were tested in triplicate. Data was analyzed manually looking at individual video frames. The number of flies that climbed past the target mark on the vial after 90 seconds was recorded.

### Bang sensitivity assay

To analyze seizure phenotypes, bang-sensitivity assays were performed essentially as previously described (Kroll and Tanouye, 2013; Song et al., 2008). 100 female flies per genotype were collected within 24 hours after eclosion under CO_2_ and moved to a fresh vial with media and allowed to recover overnight. For testing, flies were transferred into an empty vial and vortexed at maximum speed for 10 seconds. The bang-sensitive phenotype was scored as the number of flies that did not experience paralysis or seizure lasting more than 20 seconds. Flies for each genotype were collected and tested over a period of several days and the results of each experiment pooled.

### Statistics

All data was recorded in Microsoft Excel and processed using Prism 9 (Graphpad). Results were statistically significant at p < 0.05. Results shown throughout the study are mean ± SEM. n.s. = not significant, * p < 0.05, ** p < 0.01, *** p < 0.001, and **** p < 0.0001. Statistical tests used for each experiment are found in respective figure legends.

## RESULTS

### Generation of new *scat* alleles

To examine *scat* function in adult *Drosophila*, we first generated new alleles by mobilizing the P-element insertion located in the 5’ end of the *scat^1^* allele (Fig. 1A). *scat^1^* is thought to be a protein null that causes male sterility and defects and the larval NMJ (Castrillon et al., 1993; Fari et al., 2016; Patel *et al.*, 2020). We isolated two partial deletions of *scat* (*scat*^Δ*244*^ and *scat*^Δ*312*^). The *scat*^Δ*312*^ allele starts at the P-element insertion and deletes 1571 bp of downstream sequence (Fig. 1A). The *scat*^Δ*244*^ allele removes the P-element and a 346 nt fragment upstream of the insertion site that spans an intron-exon junction (Fig. 1A). When homozygous, both lines produced viable adults and, as with the original *scat^1^* line, males were sterile (data not shown). Finally, a genetic control line was generated by precisely excising the P-element (*scat^329PE^).* This line rescued the male sterility phenotype observed with the *scat^1^* and mutant alleles.

**Figure 1.**
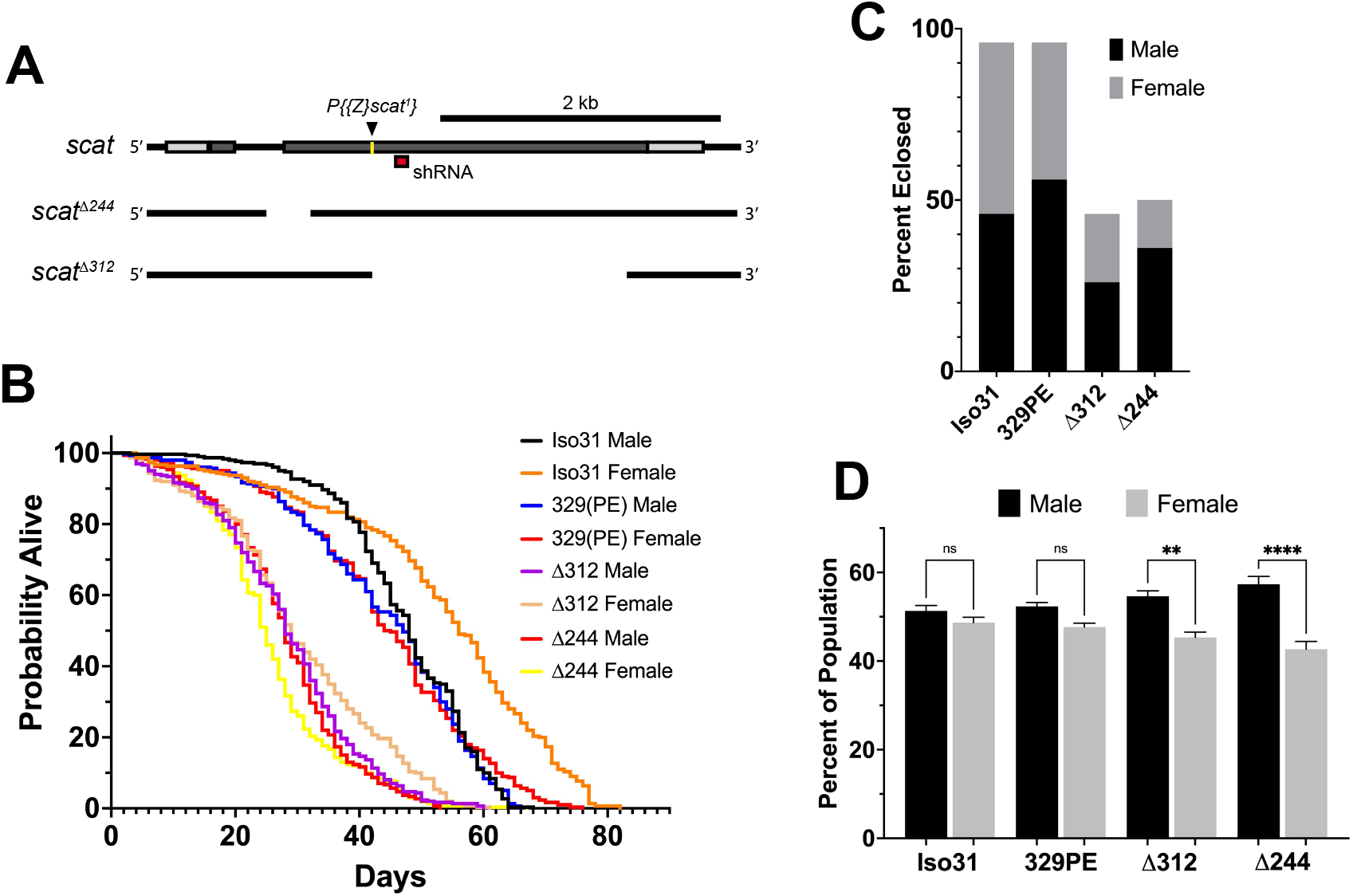
*scat* mutant adults have a shortened lifespan and a male-biased sex ratio. (A) Schematic representation of the *scat* gene showing the insertion site for the *scat^1^* P-element (arrowhead) and location of the target sequence for the *scat* shRNA (red box). The location of the deleted sequence in *scat^Δ244^* and *scat^Δ312^* are indicated by gaps. (B) Lifespan analysis of homozygous flies of the indicated genotype and separated by sex (n=300). (C) The proportion of homozygous flies of the indicated genotype that survived pupation by sex (n=50). (D) Sex ratio of flies homozygous for the indicated genotypes (n=100). Statistics: Ordinary one-way ANOVA with Holm-Sidak post-hoc analysis.

### *scat* mutant adults have a shortened lifespan and male-biased sex ratios

The wobbler mouse has a significantly shortened lifespan (Duchen and Strich, 1968). To determine if *scat* mutants also exhibited longevity defects, we conducted lifespan studies in adult flies that were homozygous for each *scat* allele. In contrast to *Iso31* controls, *scat^329PE^* females did not live longer than males suggesting there is some effect of genetic background on longevity in *scat^329PE^* females (Fig. 1B and Table 1). Thus, statistical comparisons have been made to the more genetically similar *scat^329PE^* controls. Importantly, the median lifespans of both *scat*^Δ*244*^ and *scat*^Δ*312*^ males and females were significantly reduced (Table 1). As with *scat^329PE^*, only a small (but significant) difference in lifespan was observed between *scat*^Δ*244*^ and *scat*^Δ*312*^ males and females (Table 1). Together, these data suggest that disruption of *scat* results in a reduced lifespan.

**Table 1.** Lifespan analysis of *scat* mutants.

| Table 1. Lifespan analysis of *scat* mutants |  |  |  |  |
| --- | --- | --- | --- | --- |
|  | Median life (days) | Difference vs. sex-matched *scat*<sup>329PE</sup> (%) | Significance vs. sex-matched *scat*<sup>329PE</sup> | Significance between sexes |
| *Iso31* male | 48 | 102 | n.s. | p < 0.0001 |
| *Iso31* female | 56 | 126 | p < 0.0001 |  |
| *scat*<sup>329PE</sup> male | 47 | - | - | n.s. |
| *scat*<sup>329PE</sup> female | 44.5 | - | - |  |
| *scat*<sup>Δ244</sup> male | 28 | 58 | p < 0.0001 | p < 0.05 |
| *scat*<sup>Δ244</sup> female | 25 | 56 | p < 0.0001 |  |
| *scat*<sup>Δ312</sup> male | 28 | 58 | p < 0.0001 | P < 0.01 |
| *scat*<sup>Δ312</sup> female | 29 | 65 | p < 0.0001 |  |

The *scat^1^* allele has been shown to be semi-lethal prior to eclosion (Castrillon *et al.*, 1993). This contrasts with *Vps54* loss-of-function in mice which causes embryonic lethality (Schmitt-John et al., 2005). Thus, we next determined whether *scat*^Δ*244*^ and *scat*^Δ*312*^ caused pupal lethality. Only 50% of *scat*^Δ*244*^ and 46% of *scat*^Δ*312*^ pupae survived to eclosion compared to 96% for both *Iso31* and *scat^329PE^* (Fig. 1C). These data suggest that *scat* loss-of-function causes significant lethality at the pupal stage.

During these experiments, we also noticed that there were more surviving male then female adults (Fig. 1C). To analyze this phenotype more closely, we quantified the proportions of eclosing adult flies from each genotype. While the proportion of male to female flies in *Iso31* and *scat^329PE^* controls was roughly 1:1, we observed a statistically significant male bias in *scat*^Δ*244*^ and *scat*^Δ*312*^ flies (Fig. 1D). These results further suggest that adult female flies may be disproportionally affected by loss of *scat* expression.

### *scat*^Δ*244*^ mutant females have a significantly reduced body size

The wobbler mouse has a significantly reduced body size relative to unaffected littermates (Duchen and Strich, 1968). To determine if a similar phenotype was observed in *scat* mutants, we imaged young adult flies from each genotype to examine body area and length (Fig. 2A). Quantification of body area revealed a significant decrease in body size in *scat*^Δ*244*^ females when compared to *scat^329PE^* controls (Fig. 2B). No other comparisons were statistically significant. Similarly, the analysis of body length along the longest line drawn from the rostral to caudal ends revealed a decrease in *scat*^Δ*244*^ females trending towards significance (Fig. S1A; p = 0.0537). Collectively, these data suggest that disruption of *scat* in adult females (but not males) results in decreased in body size.

**Figure 2.**
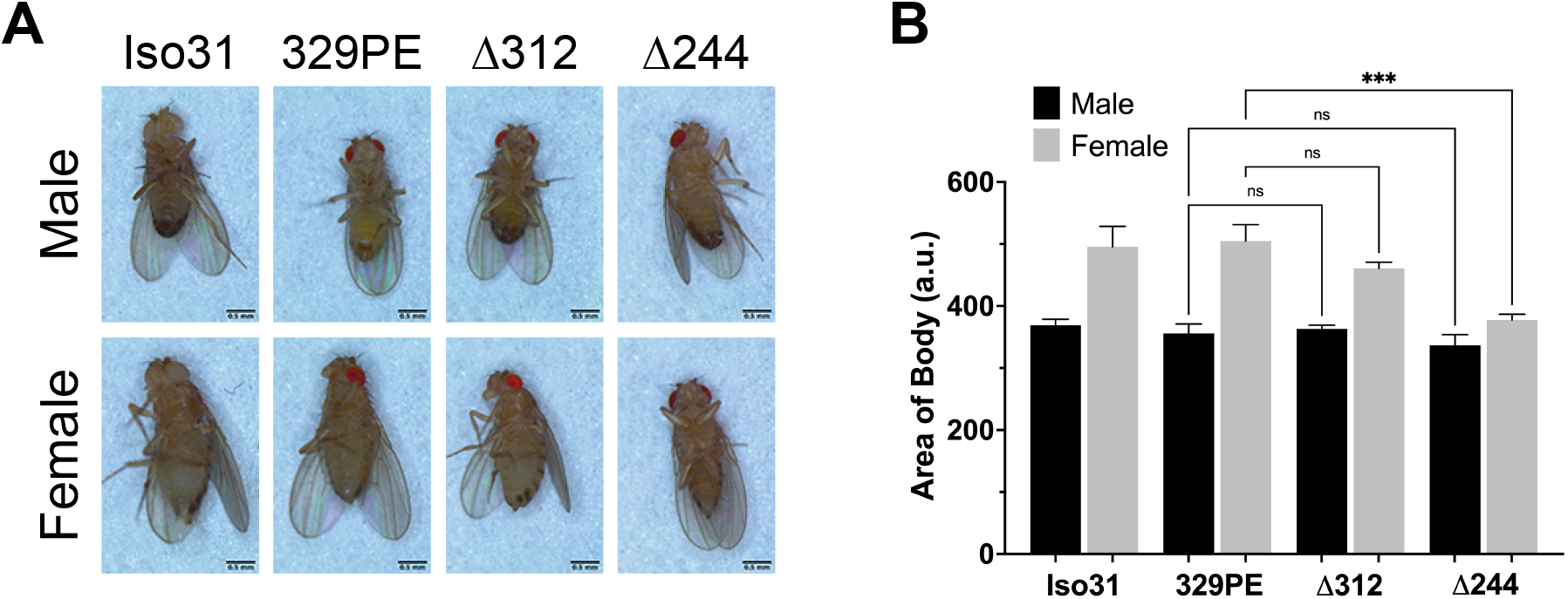
*scat* mutant females have a reduced body size. (A) Images of representative adult flies of the indicated sex and genotype. Scale bar = 0.5 mm. (B) Quantification of body area of adult flies of the indicated sex and genotype (n = 5). Statistics: Ordinary one-way ANOVA with Holm-Sidak post-hoc analysis.

### *scat* mutant females have neurological and age-progressive locomotor defects

In subsequent experiments, we focused on *scat* mutant females because they exhibit the strongest phenotypes in lifespan, viability, and body size experiments. “Bang sensitive” (bs) behavioral mutants are a means to study tonic-clonic seizures in humans (Song and Tanouye, 2008). Bang sensitivity is a phenotype where affected flies are briefly paralyzed and seize upon receiving a short mechanical shock or “bang” (Benzer, 1971; Ganetzky and Wu, 1982). Many bs mutations are in genes associated with mitochondrial function, shorted lifespan, and age-related neurodegenerative disease (Reynolds, 2018). Each of these phenotypes have also been linked to the wobbler mouse (Duchen and Strich, 1968; Santoro et al., 2004). We conducted experiments to determine if disruption of *scat* caused bs phenotypes. We found that 2-day old *scat*^Δ*244*^ and *scat*^Δ*312*^ mutant females exhibited robust bs phenotypes (Fig. 3A). This suggests that *scat* mutants are more prone to seizures, although it does not explain how this mechanistically occurs.

**Figure 3.**
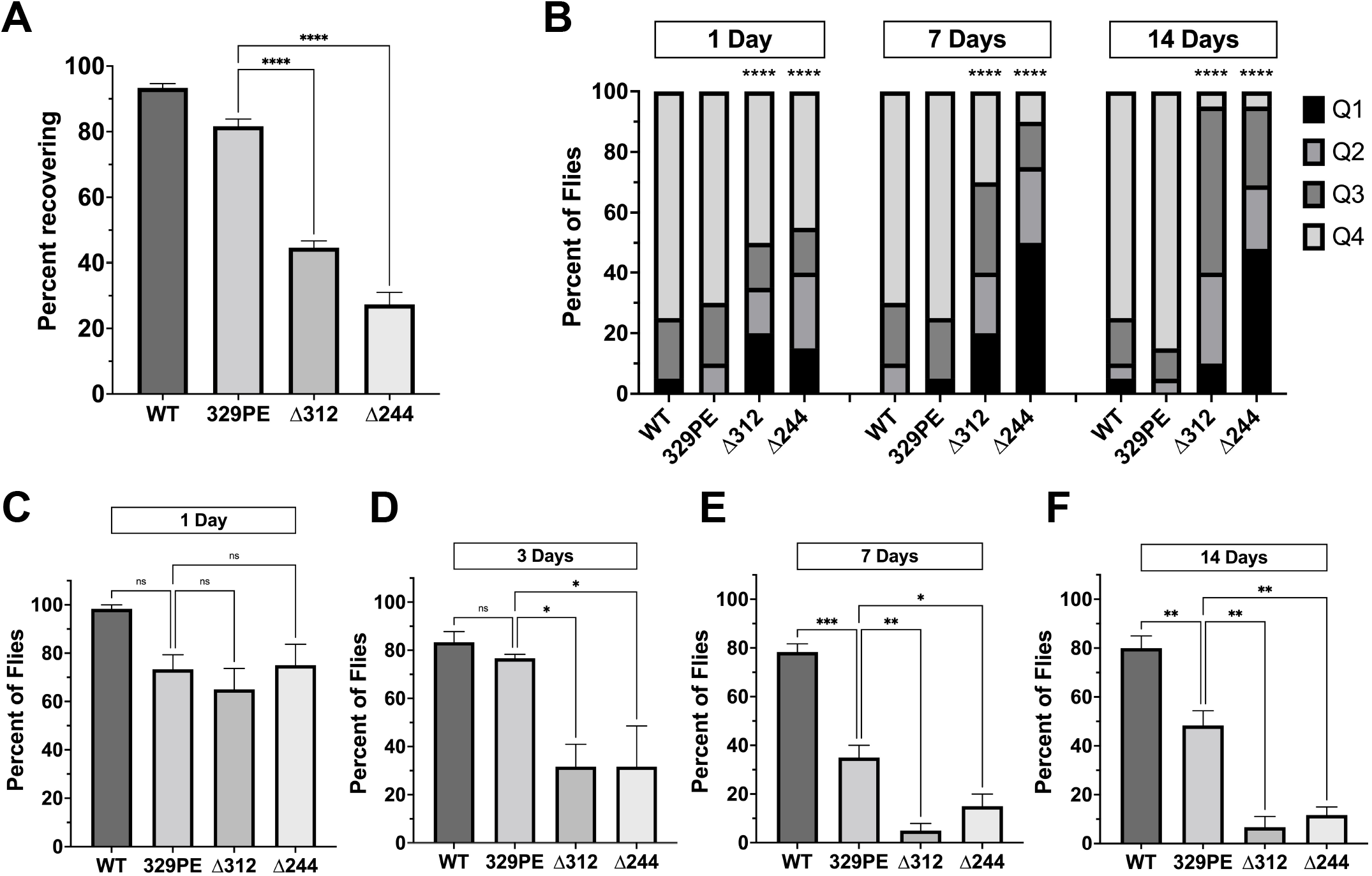
*scat* mutant females have neurological and age-progressive locomotor defects. (A) Quantification of the number of female flies of the indicated genotypes that recovered within 20 seconds following vortexing in bang sensitivity assays (n=100). (B) Quantification of the number of flies that landed in each quadrant in spontaneous flight assays. Q1 indicates the highest quadrant and Q4 is the lowest (n = 20). (C-F) Quantification of the number of female flies of the indicated genotypes able to cross the 1 cm threshold after climbing for 30 seconds (n = 20) at (C) 1 day, (D) 3 days, (E) 7 days, and (F) 14 days after eclosion. Statistics: (A and C-F) Ordinary one-way ANOVA with Holm-Sidak post-hoc analysis. (B) Chi square analysis compared to 329PE control.

Another phenotype associated with the wobbler mouse is an age-progressive motor defect caused by MN degeneration and muscle atrophy (Duchen and Strich, 1968). Therefore, we next determined whether female *scat* mutants had analogous locomotor dysfunction. To identify locomotor defects associated with primary muscle groups, we performed a spontaneous flight assay to assess the function of flight muscles. We observed a significant decrease in flight ability in both *scat*^Δ*244*^ and *scat*^Δ*312*^ as early as one day following eclosion (Fig. 3B). Moreover, while *Iso31* and *scat^329PE^* showed little change at two weeks of age, flight ability in *scat*^Δ*244*^ and *scat*^Δ*312*^ progressively worsened over time (Fig. 3B). To further examine adult locomotor ability, we performed a climbing assay. One major benefit of this approach is that it allowed us to examine the same group of females as they aged. While climbing defects were not observed in *scat* mutants at 1 day of age, significant differences between both *scat* mutants and controls were observed at 3 days and this became progressively worse at 7 days post-eclosion (Fig. 3C-E). Interestingly, the *scat^329PE^* control did not climb as well as *Iso31* controls after 7 days, again suggesting that there may be a genetic background effect (Fig. 3E-F). Together, these data suggest that *scat* mutants have age-progressive locomotor defects.

### *scat* mutant females have reduced size and degeneration of longitudinal muscles

Age-related flight and climbing defects observed in *scat* mutants suggest that there may be muscle dysfunction or degeneration. To address this question, we conducted a histological analysis of the major thoracic muscle in female flies at 1 and 7 days after eclosion. Light micrographs of histological sections revealed that the size and organization of the six bilaterally paired dorsal longitudinal muscles were significantly smaller in *scat* mutants (Fig. 4A-C). In *scat*^Δ*312*^ females, muscle degeneration was often observed in some longitudinal muscles (Fig. 4A). In contrast, this phenotype was never observed in either *Iso31* or *scat^329PE^* controls. The reduction in muscle area became significantly more pronounced in older flies (Fig. 4C). Interestingly, this “compacted muscle” phenotype is most like those observed in *Drosophila* models for myotonic dystrophy type 1 (DM1) (Bargiela et al., 2015; Garcia-Lopez et al., 2008).

**Figure 4.**
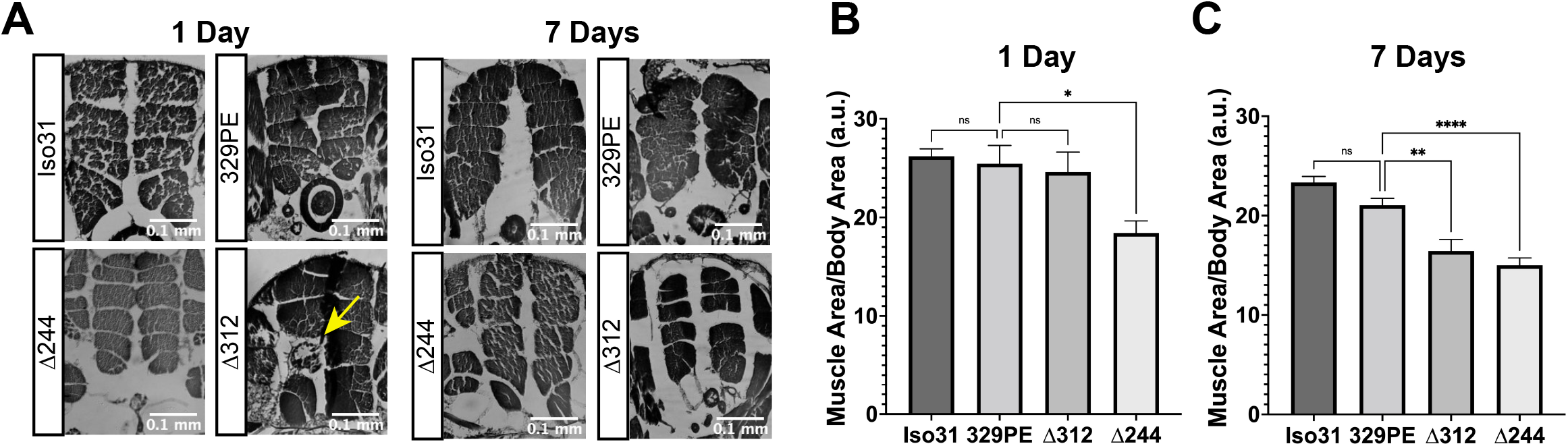
*scat* mutant females have defects in muscle size and signs of atrophy. (A) Representative H&E stained thoracic muscle sections of female flies of the indicated age and genotypes. Sections were obtained in the same region of the thorax and oriented so that the dorsal axis is up. Yellow arrows indicate muscle with signs of atrophy. (B) and (C) Quantification of the muscle area for female flies of the indicated age and genotype (n= 11-20). Statistics: Ordinary one-way ANOVA with Holm-Sidak post-hoc analysis.

### *scat* interacts genetically with *Rab11* to control locomotion and muscle atrophy

The GARP complex interacts with and regulates the tethering of both early and late endosomes at the TGN (Conboy and Cyert, 2000; Conibear et al., 2003; Conibear and Stevens, 2000; Siniossoglou and Pelham, 2002). Early and late endosomes are defined and regulated by the small GTPases, Rab5 and Rab7 (respectively). We have previously shown that *scat* interacts genetically with *rab7* (but not with *rab5* or *rab11*) in MNs to regulate synaptic integrity and development at the MNJ in fly larvae (Patel *et al.*, 2020). We hypothesized that this interaction may persist into adulthood. For this analysis, we used an inducible transgenic *scat* short hairpin RNAi line (*UAS-Scat^TRiP^*) and lines that drive the expression of either wild-type or dominant negative (DN) forms of Rab proteins (*UAS-Rab^WT^* and *UAS-Rab^DN^*). The *scat* RNAi and *rab* DN constructs were used so that we could specifically disrupt expression (or function) of both genes only in MNs.

We first determined if *scat* interacted genetically with *rab5, rab7*, and *rab11* to regulate age-progressive locomotion in climbing assays. MN-specific overexpression of Rab proteins paired with knockdown of *scat* had no effect on climbing (Fig. S2A-B). In contrast, disruption of only *rab11* paired with *scat* RNAi significantly reduced climbing ability of female flies at 7 days post-eclosion (Fig. 5B). This was still significant even though disruption of *rab11* in controls had a negative effect (Fig. 5A-B). Surprisingly, disruption of both *scat* and *rab7* had no effect on climbing ability after 7 days (Fig. 5B). To determine the nature of the locomotor defects, we also examined the major thoracic muscles in histological sections. As with climbing, we observed no significant morphological differences when *scat* knockdown was paired with the overexpression of wild-type *rab7* or *rab11* (Fig. S2C). Strikingly, we saw significant muscle atrophy in most females when both *scat* and *rab11* were disrupted in MNs (Fig. 5C). Together, this suggests that *scat* interacts genetically with *rab11* and not *rab7* in adult MNs.

**Figure 5.**
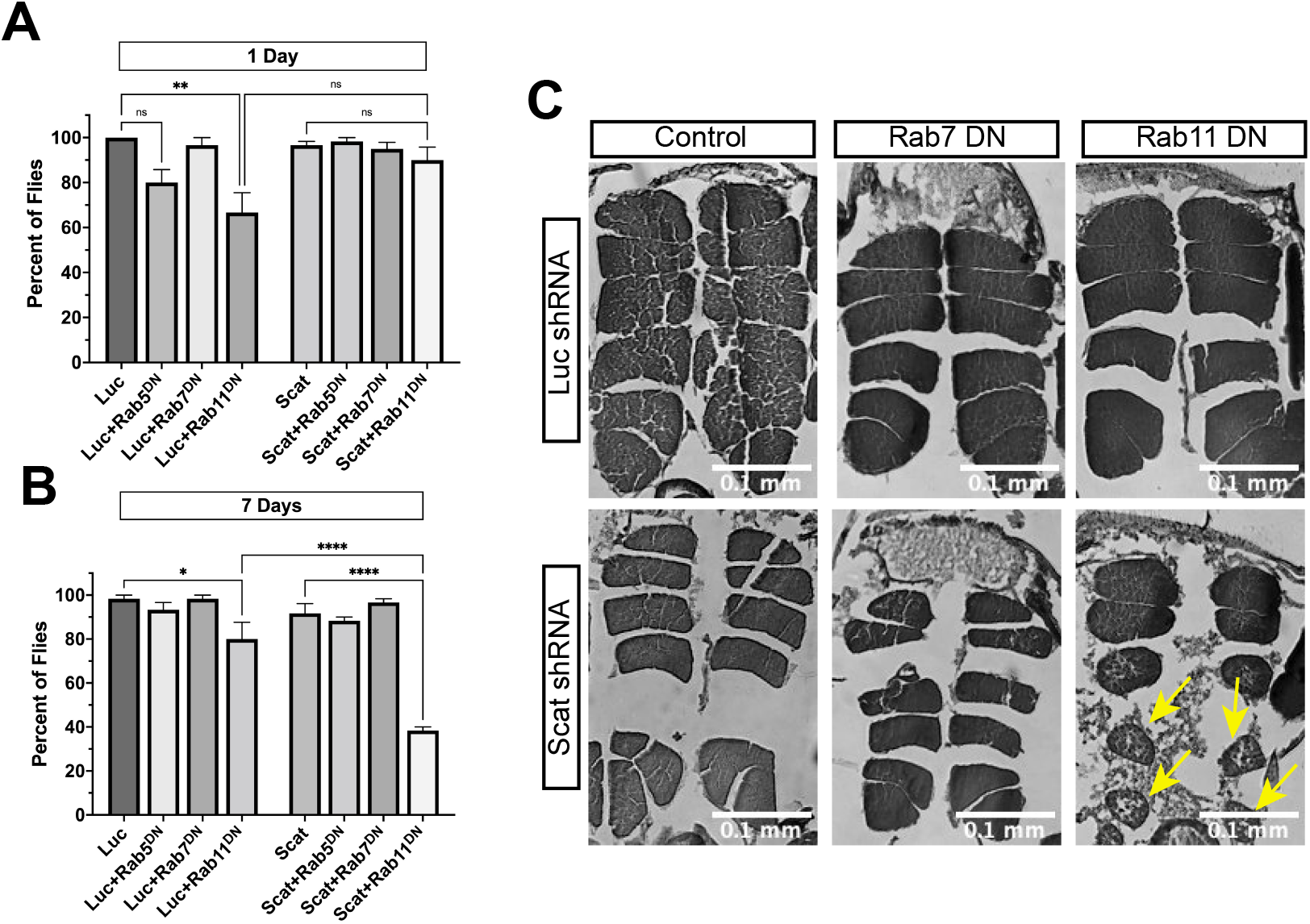
*scat* interacts genetically with *rab11* to control locomotion and muscle integrity. (A) and (B) Quantification of the number of female flies of the indicated age and genotypes able to cross the 1 cm threshold after climbing for 30 seconds (n = 20). Here, expression of either a control or *scat* shRNA transgene (*UAS-Luc^shRNA^* and *UAS-scat^shRNA^*) and dominant negative Rab transgene (*UAS-rab^DN^)* was driven in MNs by *C380-Gal4*. (C) Representative H&E stained thoracic muscle sections of female flies of the indicated age and genotypes. Sections were obtained in the same region of the thorax and oriented so that the dorsal axis is up. Yellow arrows indicate muscle with signs of atrophy. Statistics: Ordinary one-way ANOVA with Holm-Sidak post-hoc analysis.

## DISCUSSION

One of the primary objectives of this study was to determine whether disruption of *scat* expression in adult flies caused phenotypes like those observed in the wobbler mouse. It has previously been shown that *scat* mutants share spermatogenesis defects caused by Golgi dysfunction with the wobbler mouse (Castrillon *et al.*, 1993; Fari *et al.*, 2016). Here, we find that *scat* mutants share additional phenotypes including locomotor defects, decreased body size, muscle atrophy, and a shortened lifespan (Duchen and Strich, 1968). Importantly, these phenotypes are also consistent with degenerative human MN diseases, most notably ALS (Bruijn et al., 2004). While not all wobbler phenotypes could be examined here, our data suggests that *scat* loss- or MN-specific reduction-of-function may serve as a new model to study progressive MN disease.

There are some notable differences between *scat* mutant flies and the wobbler mouse. First, loss-of-function of Vps54 in mice causes embryonic lethality characterized by the underdevelopment of cardiac muscle and motor neurons (Schmitt-John *et al.*, 2005). In contrast, loss of *scat* expression in flies causes only partial lethality, primarily in females. Second, the loss of MNs in the wobbler mouse leads to muscle spasticity and not seizures, as we observe in flies (Duchen and Strich, 1968). However, mutations in Vps53 have been linked to the seizure phenotypes observed in pontocerebellar hypoplasia type 2E (PCH2E) suggesting that seizures may be associated with disruption of the GARP complex (Feinstein et al., 2014). Finally, there is no evidence of phenotypic sexual dimorphism in the wobbler mouse. Moreover, while sex has been reported to be a significant factor influencing ALS development, males have been found to be more susceptible than females (Trojsi et al., 2020). It is likely that our results are due to differences between *Drosophila* and mammalian neurophysiology. For example, there is a significant amount of evidence suggesting that female-specific steroid hormones like estrogen, that are lacking in flies, have neuroprotective properties (Zarate et al., 2017).

While we do not directly show that disruption of *scat* causes MN phenotypes, we provide evidence in support of this hypothesis. We find that knockdown of *scat* in the pre-synaptic MN paired with disruption of Rab11 activity significantly reduces locomotor ability of females and causes atrophy in the postsynaptic muscle (Fig. 5A-C). Similarly, we have previously shown that presynaptic knockdown of *scat* combined with disruption of *rab7* in larval MNs disrupts the integrity of postsynaptic densities at the NMJ via an unknown mechanism (Patel *et al.*, 2020). We speculate that neuromuscular dysfunction starts to occur in larvae and begins to manifest as muscle atrophy during metamorphosis and early adulthood (Kuleesha et al., 2016). Muscle atrophy is commonly associated with neurodegenerative disorders involved in the peripheral nervous system such as ALS, PD, multiple sclerosis (MS), and Charcot-Marie-Tooth disease (CMT) (Allen et al., 2008; Dyck and Lambert, 1968; Harding and Thomas, 1980; Leger et al., 2006; Peker et al., 2018). As MNs progressively denervate myofibrils, muscle atrophy occurs, preceded by a decrease in sarcolemma permeability. This most commonly manifests in neuromuscular disorders as muscle weakness and loss of muscle mass (Cisterna et al., 2014).

Why is there a transition to *rab11* in adults? Rab11 mediates endosome recycling to the TGN and plasma membrane and regulates the function of recycling endosomes (REs) (Kelly et al., 2012). While Vps54 does not interact with REs directly, they are important components involved in vesicular recycling and they a play critical role in axon development, axon pathfinding, synaptic vesicle recycling, and synaptic plasticity (Rozes-Salvador et al., 2020). Levels of *rab11* are downregulated in many neurodegenerative diseases including ALS (Zhang et al., 2020). Finally, Rab11 has neuroprotective effects. For example, the overexpression of *rab11* in neurons rescues synaptic and locomotor defects in a *Drosophila* model for Huntington’s disease (HD) (Steinert et al., 2012).

In summary, we have provided data suggesting that *scat* loss-of-function flies share phenotypes that are characteristic of the wobbler mouse. Based on this, we propose that this *Drosophila* model, or “wobbler fly”, may serve a new tool to study the mechanisms that underly progressive MN disease in humans. It would be interesting at this point to determine if *scat* loss-of-function causes degeneration in adult MNs. Regardless, we can now leverage the power of *Drosophila* genetics and use this model to identify novel modifiers of the *scat* locomotor phenotypes followed by detailed characterization of genetic interactors. We have already demonstrated the utility of this model on a focused scale by our analysis of interaction with the *rab* genes.

## AUTHOR CONTRIBUTIONS

EW, ES, and SB designed the experiments. EW and ES performed the experiments. EW, ES, and SB analyzed the data. EW wrote the first draft of the manuscript. All authors contributed to manuscript revision, read, and approved the submitted version.

## ACKNOWLEDGEMENTS

Some fly strains were obtained from the Bloomington *Drosophila* Stock Center, which is funded by National Institutes of Health grant P40OD018537. We thank S. Van Engelenburg for assistance with developing the fly collar used for histological section. We also thank P. Patel for initial assistance generating *scat* deletion lines. We thank members of the Barbee lab for providing useful discussion and critical comments.

