## Supplemental Figures for "Vps54 regulates lifespan and locomotor behavior in adult *Drosophila melanogaster*"

### SUPPLEMENTARY FIGURE LEGENDS

**Figure S1. *scat* mutant females do not have a reduced body length.** (A) Quantification of body length of adult flies of the indicated sex and genotype (n=5). Statistics: Ordinary one-way ANOVA with Holm-Sidak post-hoc analysis.

**Figure S2. Knockdown of Scat and overexpression of wild-type Rabs has no effect on locomotion or muscle integrity.** (A) and (B) Quantification of the number of female flies of the indicated age and genotypes able to cross the 1 cm threshold after climbing for 30 seconds (n = 20). Here, expression of either a control or *scat* shRNA transgene (*UAS-Luc*$^{shRNA}$ and *UAS-scat*$^{shRNA}$) and wild-type Rab transgene (*UAS-rab*$^{DN}$*)* was driven by *C380-Gal4*. (C) Representative H&E stained thoracic muscle sections of female flies of the indicated age and genotypes. Sections were obtained in the same region of the thorax and oriented so that the dorsal axis is up. Yellow arrows indicate muscle with signs of atrophy. Statistics: Ordinary one-way ANOVA with Holm-Sidak post-hoc analysis.

Figure S1. *scat* mutant adult females do not have a reduced body length

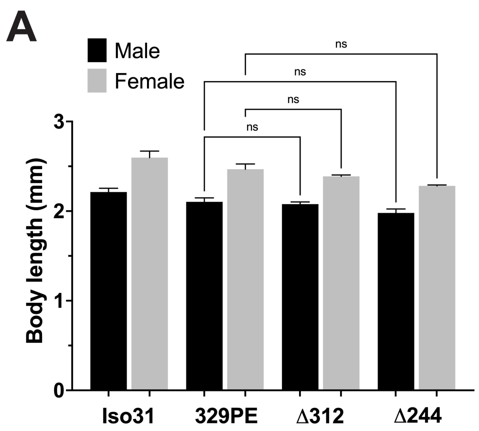

Figure S2. Overexpression of Scat and wild-type Rabs has no effect on locomotion or muscle integrity

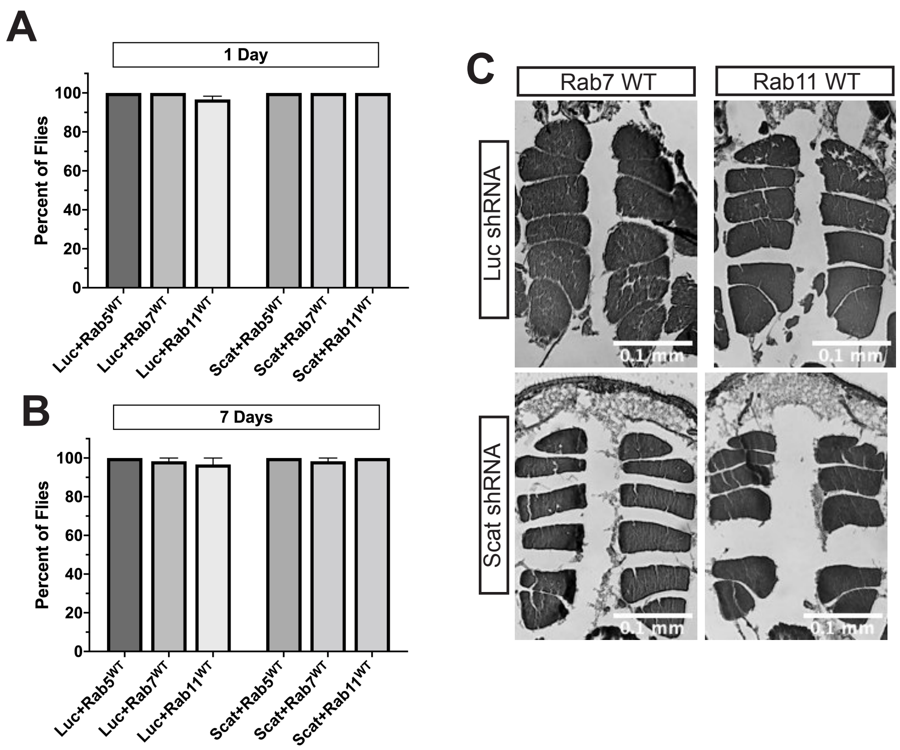
